# Molecular engineering improves antigen quality and enables integrated manufacturing of a trivalent subunit vaccine candidate for rotavirus

**DOI:** 10.1101/2020.11.20.391532

**Authors:** Neil C. Dalvie, Joseph R. Brady, Laura E. Crowell, Mary Kate Tracey, Andrew M. Biedermann, Kawaljit Kaur, John M. Hickey, D. Lee Kristensen, Alexandra Bonnyman, Sergio A. Rodriguez-Aponte, Charles A. Whittaker, Marina Bok, Celina Vega, Tarit Mukhopadhyay, Sangeeta B. Joshi, David B. Volkin, Viviana Parreño, Kerry R. Love, J. Christopher Love

## Abstract

**Background:** Vaccines comprising recombinant subunit proteins are well-suited to low-cost and high-volume production for global use. The design of manufacturing processes to produce subunit vaccines depends, however, on the inherent biophysical traits presented by an individual antigen of interest. New candidate antigens typically require developing custom processes for each one and may require unique steps to ensure sufficient yields without product-related variants.

**Results:** We describe a holistic approach for the molecular design of recombinant protein antigens—considering both their manufacturability and antigenicity—informed by bioinformatic analyses such as RNA-seq, ribosome profiling, and sequence-based prediction tools. We demonstrate this approach by engineering the product sequences of a trivalent non-replicating rotavirus vaccine (NRRV) candidate to improve titers and mitigate product variants caused by *N*-terminal truncation, hypermannosylation, and aggregation. The three engineered NRRV antigens retained their original antigenicity and immunogenicity, while their improved manufacturability enabled concomitant production and purification of all three serotypes in a single, end-to-end perfusion-based process using the biotechnical yeast *Komagataella phaffii*.

**Conclusions:** This study demonstrates that molecular engineering of subunit antigens using advanced genomic methods can facilitate their manufacturing in continuous production. Such capabilities have potential to lower the cost and volumetric requirements in manufacturing vaccines based on recombinant protein subunits.

## Background

The global demand for vaccines is on pace to exceed current capacities for manufacturing of both existing and new vaccines [1]. Increasing instances of epidemics and emerging diseases, such as COVID-19, also require constant development of new vaccines [2–4]. Despite this need, both the cost of development and the risk of failure for any given vaccine candidate are high [5]. New approaches to facilitate efficient development and manufacturing of vaccines are essential to address current and future global needs.

Subunit vaccines are increasingly prevalent due to their safety and efficacy [6–8], Examples include recombinant subunits for influenza and varicella (shingles) as well as viruslike particles (VLPs) used for hepatitis B (HBV) and human papillomavirus (HPV) [9,10]. Despite recent research on improving designs for subunit vaccines [11,12], antigens are primarily selected first for their ability to invoke appropriate immunogenic responses [13]. Further consideration of the suitability of an antigen for objectives like low-cost production during translational development could facilitate timely translation of new designs and ultimately improved global access for these products.

In contrast to subunit vaccines, the development and manufacturing of new monoclonal antibodies (mAbs) as biopharmaceuticals is highly efficient. This advantage is realized from similar processes for production for each new molecule [14]. MAbs are often amenable to efficient production using mammalian host cells, owing to high titers of secreted protein and standardized purification processes allowing for streamlined recovery. Without these features, non-mAbs require bespoke processes to produce and present distinct technical challenges. For example, many subunit vaccines are produced in intracellular compartments, like inclusion bodies in bacterial cells, which must be recovered by cellular lysis, multiple stages of chromatography, and often solubilization or refolding to achieve the desired conformation for immunogenicity. Reducing the number of steps in production, along with increasing the volumetric productivity for a given antigen, could maximize efficiency—and thus lower costs, of manufacturing [15,16].

Here, we present an approach informed by host cell biology to engineer subunit vaccine candidates for efficient manufacturing while retaining their immunogenicity. We engineered a trivalent subunit vaccine candidate for rotavirus to enable an improved process for production that uses 50% fewer steps (relative to a reference process based on intracellular bacterial expression and recovery) [17]. We minimally modified the sequences of the three candidate protein antigens to promote secretion by a yeast host using a combination of genomic sequencing methods to inform specific changes for each antigen. These targeted changes improved the secreted titer and quality for each serotype antigen, addressing molecular variants affecting the quality attributes of these proteins, including aggregation, unwanted glycosylation, and *N*-terminal truncation. We show all three antigens were compatible with concomitant expression and co-purification in a single campaign. This study provides a proof-of-concept for integrated manufacturing of a multi-valent subunit vaccines while retaining quality and relative quantities of the antigens.

## Results

Rotavirus is a leading cause of mortality in children under five globally [18]. Three viral serotypes are most prevalent (P[4], P[6], and P[8]) [19]. While live viral vaccines are available for rotavirus, low-cost subunit vaccines could facilitate improved global coverage [17,20,21]. A trivalent non-replicating rotavirus vaccine (NRRV) designed at the NIH is currently advancing in clinical development. This candidate vaccine comprises three recombinant fusion-protein antigens based on VP8 truncations of the serotypes P[4], P[6], and P[8] [22]. Each VP8 truncation is genetically fused at the N-terminus to the P2 epitope of tetanus toxin (Fig. 1A). The current manufacturing strategy for these three serotypes involves fermentation in *Escherichia coli*, and recovery by multiple steps of centrifugation, cell lysis, and chromatography, followed by blending of the three antigens and formulation with adjuvant [23]. While the three serotypes have largely homologous amino acid sequences (Fig. 1B), they are different enough to present unique manufacturing challenges, making this subunit vaccine candidate an ideal case study for molecular engineering to improve its production.

**Fig. 1.**
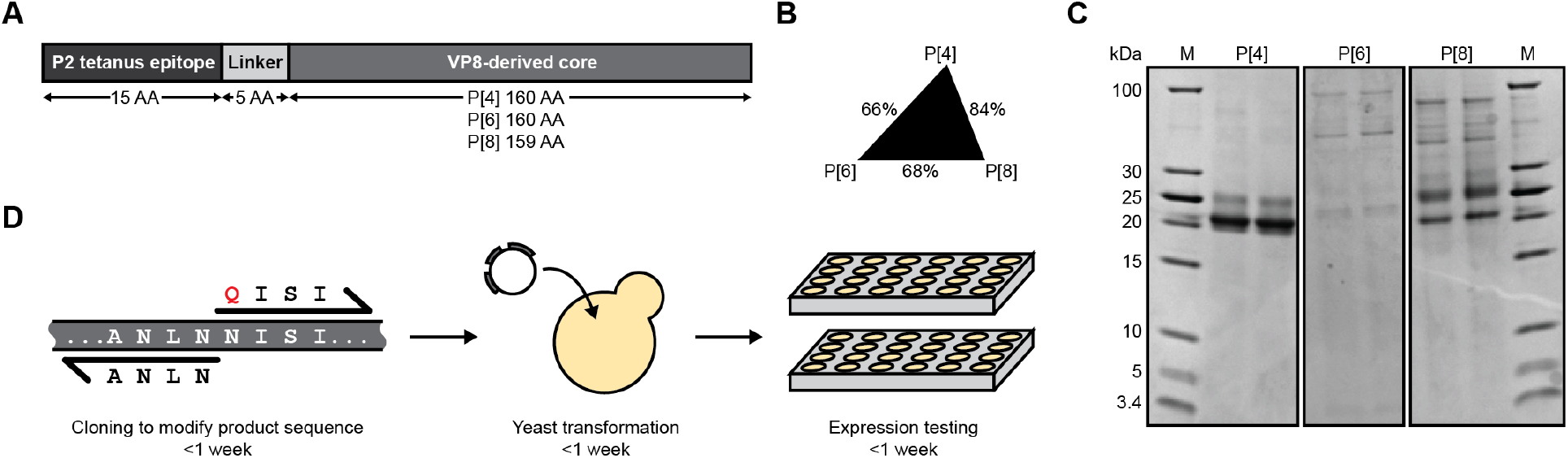
Differences in product titer and quality among three non-replicating rotavirus vaccine (NRRV) serotypes provide case studies for sequence engineering. A) Diagram of the three NRRV subunit vaccine antigens. AA = amino acid. B) Percent identity of amino acid sequences between the three subunit antigens. C) SDS-PAGE of the initial expression of each NRRV antigen. Expression from two clones is shown for each antigen. D) Diagram of a design cycle for sequence engineering.

Secretion of recombinant heterologous proteins by a host cell can simplify its recovery by reducing the number of non-chromatographic steps like centrifugation, lysis and refolding often required for intracellularly expressed proteins. Secretion-based strategies for production are also conducive for continuous operations that can increase the volumetric productivity in manufacturing [24]. For these reasons, we sought to enable secretion of the three antigens in the yeast *Komagataella phaffii*. We tested the secreted expression of each NRRV serotype independently in *K. phaffii* using the reported amino acid sequences expressed in *Escherichia coli* [23]. To adapt the sequences of each of the three antigens for this purpose, we optimized the codons of the genes for each serotype for expression in *K. phaffii*, added a signal peptide for secretion into the culture supernatant, and then assessed protein titers and quality following small-scale batch cultivation (Fig. 1C). Each serotype presented a markedly different set of molecular variants when recovered. The P[4] antigen was observed with a titer of ~50 mg/L in plate-scale culture (Fig. S1A), but exhibited a high fraction of low-molecular weight variants. The P[6] antigen was barely detected in culture supernatant or intracellularly (Fig. S1B). Secreted P[8] was extensively hypermannosylated and we detected product-related high-molecular weight variants (HMWV) (Fig. S2A-B). These variations in antigen expression and quality presented challenges in developing a robust process for manufacturing, and did not suggest that a process using common steps or operations likely would be possible. Separation of product-related variants such as those observed with P[4] and P[8] typically require additional customized steps. The poor secreted titers observed with P[6] might also require extensive development of fermentation conditions and media or intracellular expression and subsequent recovery to achieve sufficient productivity for cost-effective manufacturing.

We sought to engineer each serotype to improve its manufacturability and quality attributes, specifically to address the product-related variants identified (glycosylation, sequence variants, HMWV), without impacting the immunogenicity of the reference antigens. In our approach, we adopted a strategy similar to the guidance for biopharmaceuticals for Quality by Design [25]. For each antigen, we identified the key attributes for each that influenced their secretion by host cells and the complexity of a purification process required for recovery. Informed by these heuristics and data collected during characterization of the host cell biology experienced during protein secretion, we designed and tested a limited number of conservative sequence modifications for their impact on protein titers and observed quality attributes. Each design cycle comprised modification of an antigen sequence, generation of a strain, and evaluation of secreted protein titer and product-related variants. Iterative cycles were executed in less than three weeks in *K. phaffii* (Fig. 1D), enabling the rapid assessment of how variations in sequence motifs impacted several key molecular attributes in the secreted proteins. We describe below the approach used to modify each serotype.

### Modification of amino acid sequence eliminates product-related variants

Based on our analysis of the secreted P[8] antigen, we identified unwanted glycosylation and the formation of dimers as key molecular attributes to address; these variants can both complicate separation by chromatographic methods. *N*-linked glycosylation and dimers resulted from specific known amino acid motifs (N-X-S/T) and an unpaired cysteine, respectively. Four sites in P[8] were predicted by the NetNGlyc tool [26] to have potential for *N*-linked glycosylation during protein folding and export (Table S1). We altered the P[8] sequence, therefore, to remove the most confidently predicted glycosylation sites using a conservative, targeted change (NàQ). After elimination of the two most likely glycosylation sites (N85 and N151), no hypermannosylation was detected on P[8] by SDS-PAGE or LC-MS (Fig. 2A, Fig. S3).

**Fig. 2.**
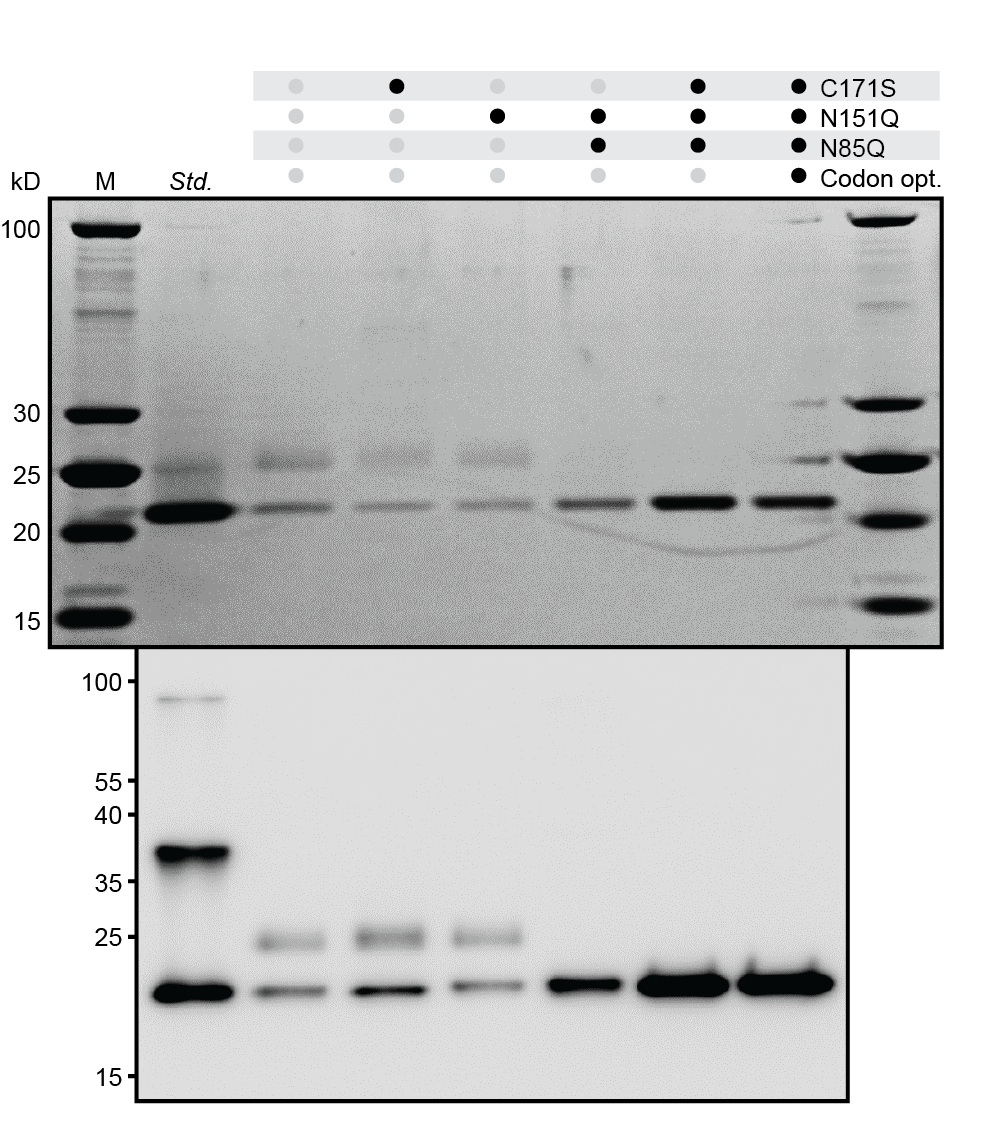
Sequence changes to remove *N*-linked glycosylation and Cys dimerization improve the quality and titer of P[8]. SDS-PAGE (top) and Western blot (bottom) of variants of P[8].

In parallel, we generated CàS modifications of the free cysteine present in the P[8] antigen (C171) and observed nearly complete elimination of HMWV (Fig. 2A). When combined, these changes at C171, N85, and N151 also increased the secreted titer of aglycosylated, monomeric P[8] from ~26 to ~87 mg/L in plate-scale batch cultivation. These data affirmed that specific, targeted changes can simultaneously improve multiple quality attributes of the antigen and the titers of secreted protein.

### Product nucleic acid sequence influences N-terminal fidelity and folding

We then applied the same principles to address the observed molecular variants in the P[4] antigen. Initial expression of P[4] revealed multiple product-related variants with lower molecular weights by SDS-PAGE; these were confirmed by intact MS as four discrete *N*-terminal truncations (Fig. 3A). Interestingly, P[8] exhibited less truncation than P[4] despite having an identical *N*-terminal amino acid sequence (the P2 tetanus epitope). We theorized, therefore, that alternative translation start sites or ribosome skipping did not induce the observed differences in truncation [27,28].

**Fig. 3.**
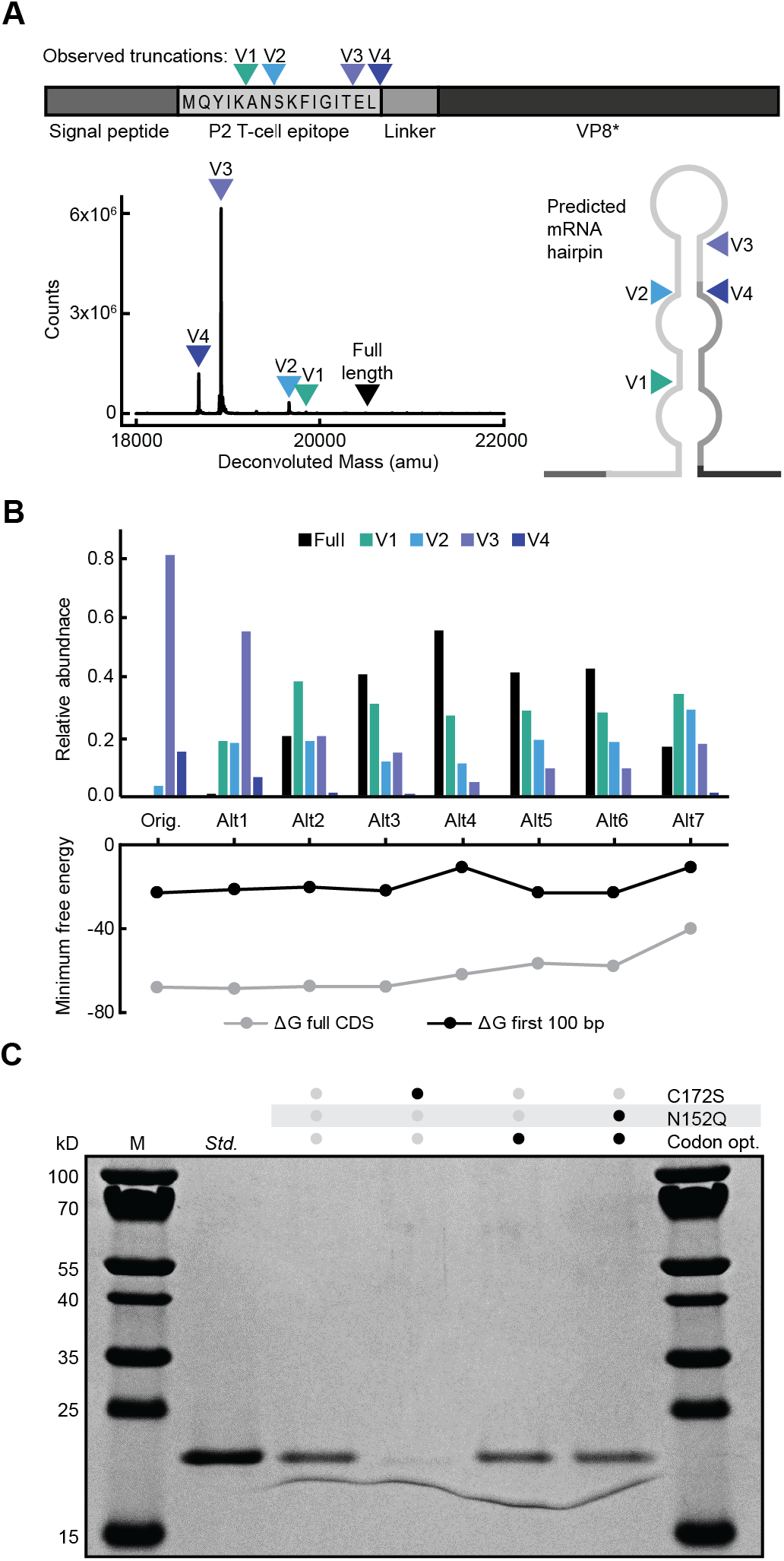
Codon optimization reduces the *N*-terminal truncation of P[4]. A) Truncation variants of the original P[4] sequence by LC-MS. The N-terminus of each truncation variant is indicated on the antigen diagram and predicted RNA hairpin structure. B) Relative abundance within the sample of truncated P[4] variants identified by LC-MS (top). Free energies of RNA hairpin formation after codon optimization for each variant (bottom). C) SDS-PAGE of variants of P[4].

Alternatively, it has been noted that undesired RNA structures, such as hairpins in nascent mRNA, can cause ribosomal stalls and truncations [29]. We hypothesized that secondary structures in the transcript could impact the variants manifest in the recovered polypeptides. Using RNAstructure [30], we identified an energetically favorable mRNA structure predicted to form near the sites of truncation in P[4] (Fig. 3A). We modified the codon usage of the P2 tetanus epitope to reduce the strength of the predicted RNA secondary structure while preserving the amino acid sequence (Fig. 3B). With increasing mitigation of the predicted RNA secondary structure from −22.8 to −10.7 kcal/mol, the fraction of full length P[4] in culture supernatant increased from 0% to nearly 60%, while the fraction of the longest truncated variant (V1) increased from 0% to nearly 30%. Overall, the percentage of the antigen that was fulllength or nearly full-length (V1) increased from 0% to nearly 90% through improved codon usage alone, confirming that RNA sequence was at least partially responsible for the observed *N*-terminal truncation.

Similar to P[8], we observed hypermannosylation in the P[4] antigen and we identified sites predicted to be glycosylated. Modification of a single *N*-linked glycosylation site (N152Q) resulted in comparable titer and product quality as observed for P[8] (Fig. 3C). Surprisingly, despite strong similarity between P[8] and P[4], removal of the free cysteine residue (C172S) drastically reduced the secreted expression of P[4] and was not included in the selected optimized sequence.

### Genome-wide host cell characterization enables alleviation of translation stall sites

During initial expression, the P[6] variant strain yielded ~8-fold lower titers of secreted protein than either the P[4] or P[8]-expressing strains, making titer a key process-related parameter to optimize for this serotype. Productive secretion of a recombinant protein requires proper mRNA transcription, translation, protein translocation and folding, and vesicular export. This complex, multi-step process makes diagnosis of a particular bottleneck a difficult task [31]. We used transcriptomics to survey differentially expressed genes in the strain expressing the P[6] antigen compared to strains expressing either P[4] and P[8]. The transcripts of P[6], P[4], and P[8] each were expressed at comparable levels by their respective strains, indicating transcription is not the rate-limiting step for P[6] (Fig. 4A). Of all gene sets that were differentially expressed in P[6]-producing cells (Table S2), cytoplasmic translation had the highest enrichment score, followed by gene sets related to cell wall integrity and DNA repair, which are reported signals of stress in *K. phaffii* (Fig. 4B) [32]. We hypothesized that disrupted translation may preclude both adequate production of P[6] and proper cell growth and maintenance. We therefore focused our efforts on identifying sequence motifs that could impact protein translation.

**Fig. 4.**
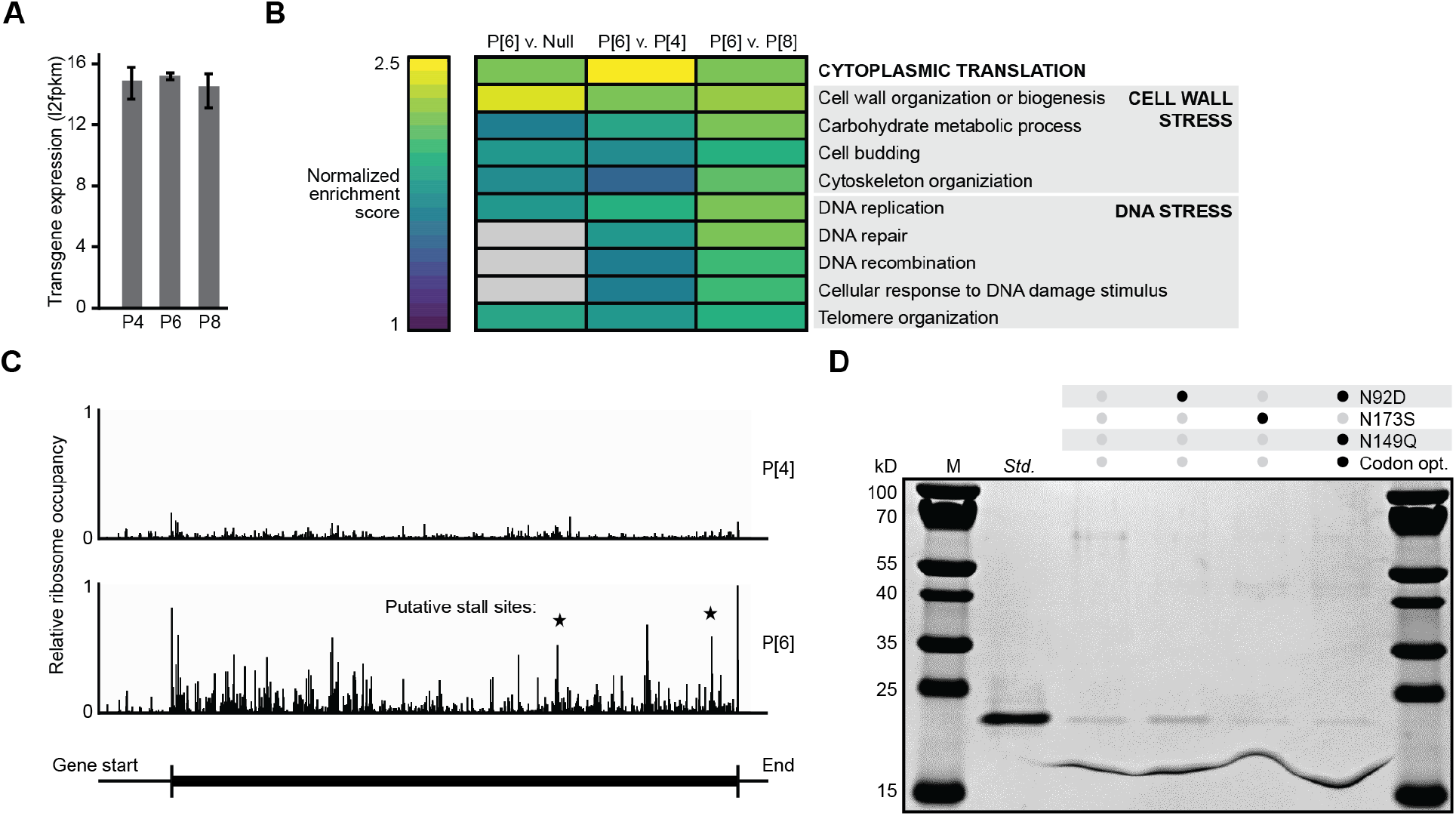
Transcriptomics and ribosome profiling reveal a translational stall site that prevents expression of P[6]. A) Expression levels of the recombinant gene in strains expressing each NRRV antigen. B) Gene set enrichment analysis of the P[6] strain compared to strains expressing P[4], P[8], or no recombinant protein (Null). Gene sets shown exhibit high enrichment scores and significant adjusted p-values (<0.05) in at least one comparison. C) Relative ribosome occupancy across the recombinant transcript in strains expressing P[4] or P[6]. The magnitude of occupancy is normalized to the occupancy on all transcripts in each strain. Putative stall sites represent peaks that are not reflected in the P[4] transcript, and are located near a sequence discrepancy between P[4] and P[6]. D) SDS-PAGE of variants of P[6].

Similar to our approach with P[4], we first examined the nucleic acid sequence of P[6] for the presence of energetically favorable RNA secondary structures using *in silico* tools for prediction. We observed a predicted RNA hairpin structure like that in P[4], but analogous mitigation of this structure did not improve production of P[6]. Upon inspection of the amino acid sequence of P[6], we noticed several highly hydrophobic amino acid motifs, which have been previously implicated in ribosomal stalling [33]. We altered amino acids near and within hydrophobic regions to reduce local hydrophobicity, but also did not observe increased expression of P[6] in any of these variants (Fig. S4).

To systematically identify candidate sites of translational stalling, we used ribosome profiling, a technique in which ribosome-associated fragments of mRNA are sequenced to determine regions of transcripts that are enriched with ribosomes [34]. We visualized ribosome occupancy at each codon of the recombinant transcript in strains expressing either P[6] or P[4] for comparison (Fig. 4C). Ribosome occupancy was determined from the number of ribosome footprints that mapped to a particular transcript, after adjusting for both the abundance of the transcript and that of footprints mapping to the native transcriptome. We observed over 4-fold higher ribosome occupancy overall on the transcript of P[6] relative to P[4], which supported our hypothesis of sequence-based translational stall sites in this serotype. We also observed sites with high ribosome occupancy at several positions of P[6] that were not similarly detected at those same positions in P[4]. Most of these positions coincided with local sequence motifs that differed in P[6] from the P[4] and P[8] consensus sequences. We nominated five of these motifs with the largest potential for ribosomal occupancy as likely sites of translational stalling and modified the amino acid sequence of each site to match that of a consensus of P[4] and P[8] sequences. Given the productive secreted expression of P[4] and P[8], we hypothesized that changing these sites in P[6] to match the P[4] and P[8] sequence could mitigate a translational stall while preserving the structure and antigenicity of the protein. One of these modifications (N92D) resulted in a ~2x improvement in the expression of P[6] (Fig. 4D). The combination of RNA-Seq and ribosome profiling, therefore, enabled us to test a reasonable number of hypotheses and increase the secreted expression of P[6] to levels readily detected by SDS-PAGE with a single substitution of an amino acid.

### Engineered subunits retain structure and function

The modifications to the primary sequences of the three subunits, though modest, could alter the structural integrity and antigenicity of the proteins. We performed bio-layer interferometry as a measure of the secondary structure of each serotype using monoclonal antibodies specific to each (Fig. 5A). All but one of the altered sequences had a minimal effect on binding affinity to the antibodies, suggesting that the modified variants retained antigenicity comparable to the reference sequences. Interestingly, the removal of the free cysteine residue of P[4] significantly altered the binding affinity of this variant, in addition to reducing expression (Fig. 3C, Fig. S1) as noted previously. This observation underscores that modifications to the amino acid sequence of a protein can interfere with the structure of a protein, and that alterations need to balance both objectives for antigenicity and manufacturability.

**Fig. 5.**
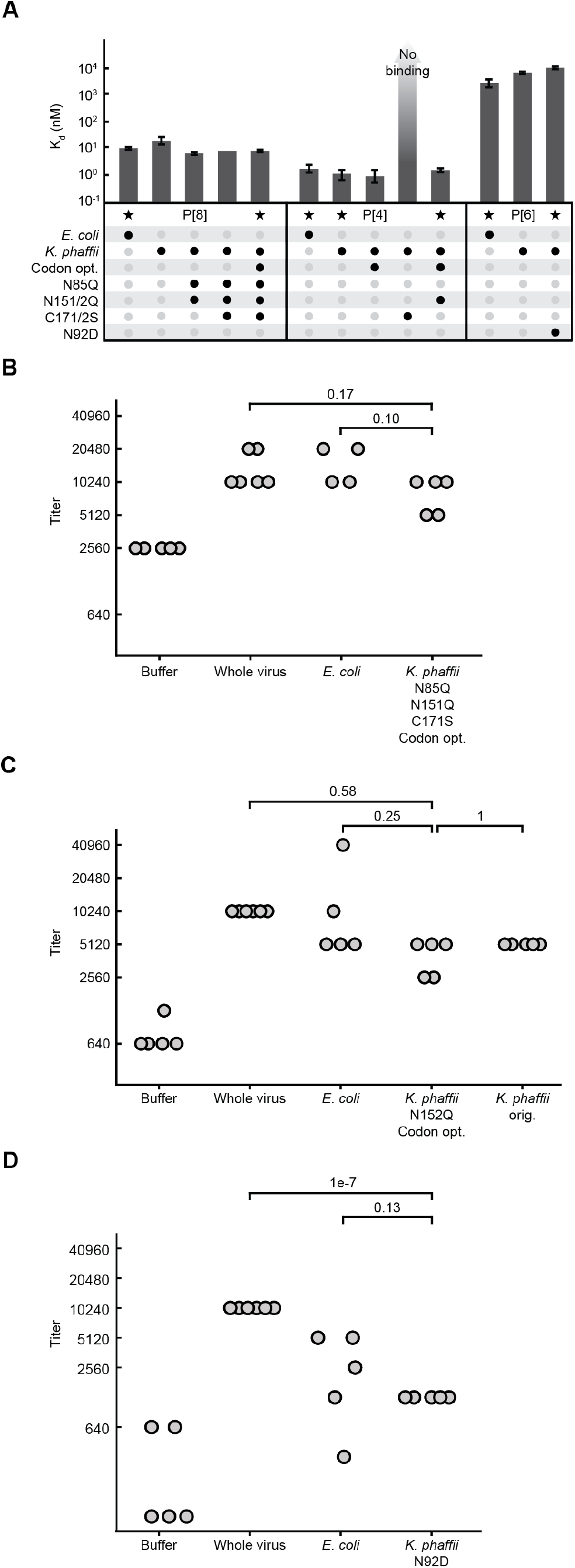
Engineered NRRV antigens produced in yeast are comparable to original antigens produced in bacteria. A) Mean binding affinity of antigen variants measured by biolayer interferometry. Variants labeled with a star were further evaluated in animals. Error bars represent the maximum and minimum binding affinity of three technical replicates. B-D) Titer of neutralizing antibodies raised against B) P[8], C) P[4], or D) P[6], measured by fluorescence-based virus neutralization assay. Each data point represents the serum of one animal, as measured 35 days after the first injection, or 7 days after the final injection. Antigens were mixed with alhydrogel before injection. Treatment groups within each serotype were compared by ANOVA, followed by *post hoc* comparison of means using Tukey’s HSD test according to best practices.[50] ANOVA results were not significant for any serotype (p>0.05), but adjusted p-values from Tukey’s test are noted.

We then evaluated whether the selected engineered antigens maintained their immunogenicity, since the modified sequences may disrupt key linear or conformational epitopes that promote immune responses. We compared the ability of the engineered antigens for each serotype to induce the production of neutralizing antibodies in guinea pigs relative to the original sequences produced in bacteria. We selected one engineered antigen per serotype having the best observed titer and quality (Fig. 5A). We produced these three engineered antigens, alongside a control having the original P[4] sequence, by secreted expression in *K. phaffii* and purified the proteins to assess their immunogenicity in adjuvanted doses. Groups of guinea pigs were vaccinated with the engineered antigens produced in *K. phaffii*, and compared directly to groups that were vaccinated with antigens produced in *E. coli* having the original sequences [20]. We measured the titers of neutralizing antibodies present in serum collected from all animals in our study after three administered doses. The measured titers of neutralizing antibodies raised in response to our engineered antigens were not significantly different from titers raised by the original antigens for all three serotypes (Fig. 5B-D). Interestingly, we observed cross neutralization among serotypes for both engineered and original antigens (Fig. S5A-C), evident by the strong neutralizing activity against P[4] virus using sera from groups vaccinated with P[6]. This cross neutralization was also evident in antigen-specific ELISAs, which confirmed general cross reactivity among the serotypes (Fig. S5D). Cross reactivity is expected given the high homologies and biophysical similarities among serotypes, and may contribute to the broadly neutralizing responses achieved by the trivalent vaccine [17]. These results suggested that the engineering changes to the three antigens implemented to improve manufacturability did not significantly alter their immunogenicity in this model.

### Reduction of product-related variants enables a common purification process

Given the overall similarity in sequence among all three antigens and the reduction in product-related variants after molecular engineering of each antigen, we hypothesized that all three antigens could be co-expressed and purified in a single cycle of production. Multi-valent subunit vaccines, in general, are strong candidates for co-purification because antigen serotypes often have similar biophysical characteristics, and integrated production could reduce the total number of manufacturing campaigns required. We previously demonstrated an integrated, continuous manufacturing system for producing biopharmaceutical proteins (InSCyT) [35]. Two core elements—microbial perfusion-based fermentation and integrated chromatographic operations [36,37]—made it possible to rapidly and simultaneously develop processes for production and recovery on this system. We first developed a single process for perfusion-based fermentation based on our prior experience producing similar molecules, and executed it to produce each antigen individually (Fig. S6A-C). Starting from processes reported previously [35], we then developed a process for purification comprising a multimodal cation exchange capture resin for the removal of host-cell proteins followed by an anion exchange resin for the removal of host-cell DNA. The same two-step process was sufficient to recover each of the three individual antigens and remove residual host-cell proteins to below 500 ppm and DNA below the limit of detection of our assay (Fig. S6D-G). Product-related variants were already sufficiently minimized by sequence engineering such that no additional purification steps were required.

### Co-expression and co-purification enable manufacturing of three antigen serotypes in a single campaign

We proceeded to produce all three antigens in a single integrated, end-to-end manufacturing process. We created a *K. phaffii* strain that expressed all three engineered antigens required for the trivalent vaccine candidate. Each transgene was introduced in series using three orthogonal selection markers, beginning with the serotype that demonstrated the lowest level of secreted expression by our host organism (P[6], P[4], then P[8]) (Fig. 6A). After each transformation of a new transgene, secreted expression of the antigen was verified by SDS-PAGE (Fig. S7A-B). We then used this strain producing all three antigens in the InSCyT system (Fig. 6B). Cells were cultivated in perfusion fermentation, and the perfusate was subsequently purified in two steps by multimodal cation exchange chromatography followed by anion exchange chromatography, as performed before for the individual antigens. Production of all three serotypes was sustained over 108 hours, and the relative ratios of each serotype were largely preserved throughout the campaign (Fig. 6C). Successful co-expression and copurification of all three serotypes were confirmed by IEF (Fig. 6D) and LC-MS (Fig. 6E). Process-related impurities including host-cell proteins and host-cell DNA were reduced in the chromatographic operations below typical regulatory guidelines (Fig. S7C-D). Overall, our approach of molecular design informed by host cell biology combined with an integrated, end-to-end manufacturing system enabled production of 200 mass-equivalent doses (limited by P[6]) of high-quality trivalent antigens in just six days on a single subliter benchtop system.

**Fig. 6.**
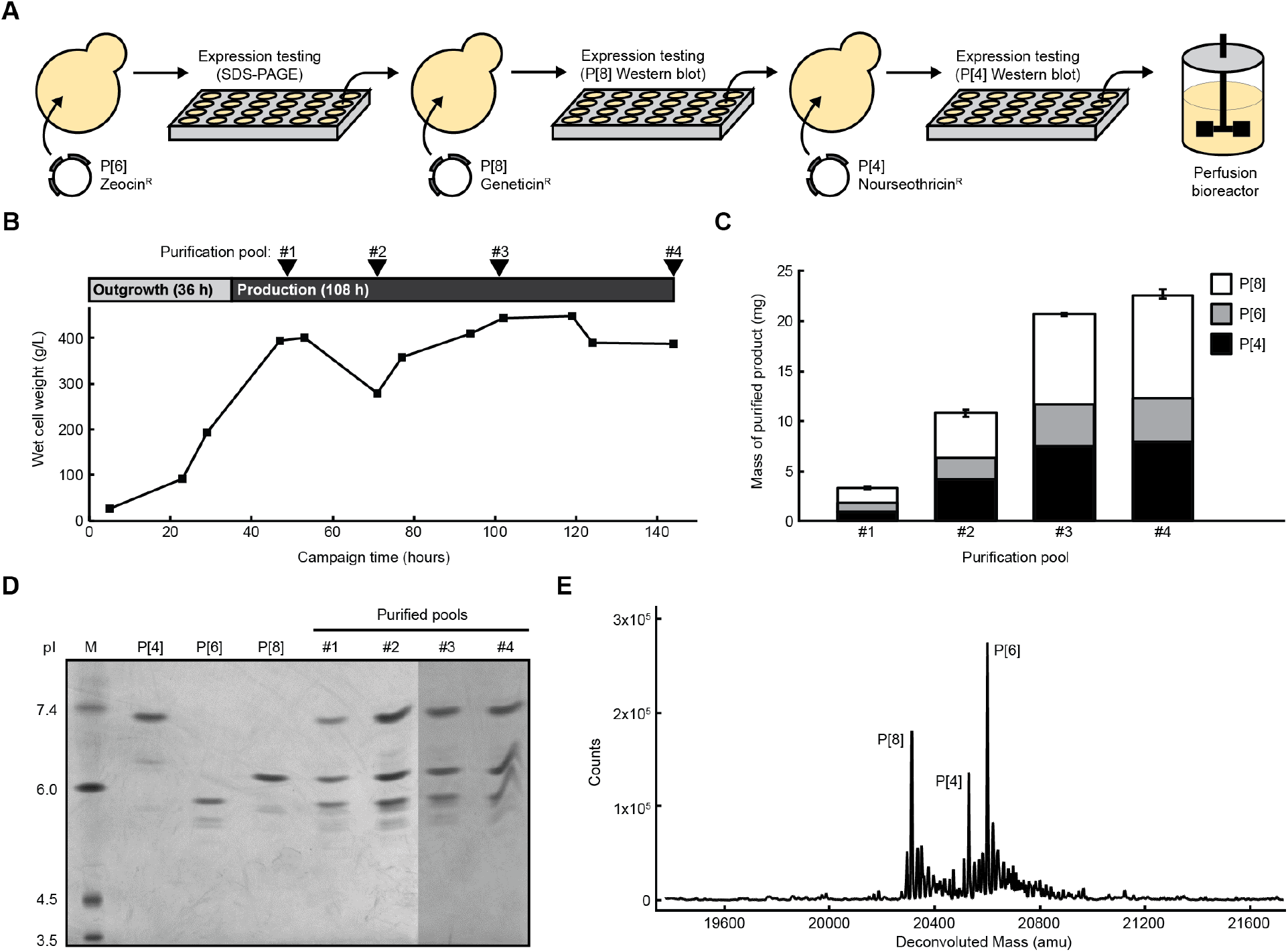
Sequence engineering enables sustained production of a trivalent NRRV vaccine from a single strain in a single manufacturing campaign. A) Construction and testing of a strain that expresses all three NRRV antigens. B) Cell density across the manufacturing campaign. Periods of outgrowth on glycerol, induction of protein expression on methanol, and points of product pooling and purification are shown. C) Yield of NRRV antigens from the campaign. Concentrations were determined by spectrophotometric absorbance at 280 nm. Ratios of antigens were determined by reverse-phase liquid chromatography. D) Isoelectric focusing gel of each purified pool of product. E) Product masses of each antigen after purification identified by LC-MS.

## Discussion

Here, we have shown that conservative targeted molecular engineering for a subunit vaccine candidate can improve product titers and quality while preserving antigenicity and immunogenicity. This approach relies on rational analysis using a combination of natural variations of the sequence and genome-scale analysis of the host for expression. This proof-of-concept study demonstrates that disparate antigen candidates can be intentionally designed to conform to a common process for manufacturing, like those operated on the InSCyT manufacturing system here. Our approach to address molecular variants impacting the quality of the antigens at the amino acid sequence level was demonstrated by simple mitigation of *N*-linked glycosylation and dimerization in P[8]. Identification of a small number of critical amino acid residues allowed rapid screening of exhaustive sequence variations, yielding a strongly expressed antigen with no major product-related variants. Free cysteine residues in model proteins have been shown to covalently bind to pharmaceutical additives such as thimerosal, a commonly used preservative in multi-dose formulations [38]. We confirmed subsequently (and described elsewhere) that removing the free cysteine residue from P[8] abrogated the formation of a mercury adduct that can limit stability of the NRRV vaccine (Sawant and Kaur, submitted; Kaur and Xiong, submitted).

To improve the product quality of the P[4] antigen, we also tested alternative codon optimization as a tool for product engineering and observed the thermodynamic stability of the P[4] mRNA structure impacted the *N*-terminal fidelity. The *N*-terminal truncations observed when expressing the original P[4] sequence are characteristic of promiscuous serine proteases; several of which are conserved in yeasts [39]. Based on our results, we hypothesized that mRNA secondary structures may slow translation and translocation, allowing increased cytoplasmic protease activity. Silent mutations have previously been demonstrated to impact product quality in biopharmaceutical manufacturing [40], and this potential mechanism merits further consideration in future studies. The connection between mRNA structure and protein truncation, nevertheless, demonstrates a key relationship through which truncated variants may be mitigated.

To increase the titer of P[6], we characterized the host cells themselves to generate a testable number of sequence changes. RNAseq revealed that translation was likely a critical cellular process inhibiting the secreted expression of P[6]. We did not identify a canonical ribosomal stall site, as such sites can be specific to the organism, and translation of exogenous genes in most biomanufacturing host organisms is not well understood [41]. Ribosome profiling as a technique, however, is agnostic to the mechanism of translational failure, and enabled us to identify a non-canonical site of translational stalling. Ribosome profiling captures data regarding all abundant transcripts in a single initial experiment, so future efforts could be focused to identify, and ultimately predict, translational preferences and stall sites that are specific to the host [42]. In this case, a modest substitution from the P[4] and P[8] sequence likely restored efficient translation of the P[6] transcript, though we were unable to identify the exact mechanism of failure.

Changes to the primary structure of a subunit vaccine, even conservative amino acid substitutions, could alter its antigenicity. Pathogens mutate T-cell epitopes or sequence-specific protease sites to facilitate evolutionary escape from antigen presentation to CD4^+^ T-cells [43]. Primary sequence changes may also alter secondary and tertiary structures that create discontinuous or conformational epitopes [44,45]. We recently demonstrated that physicochemical properties of the candidate antigens here are sensitive to single “weak spot” residues that may also impact antigenicity [46]. For these reasons, and as demonstrated here, conservative, isolated amino acid mutations are preferable to sequence-wide screening because they reduce the probability of disrupting unknown antigenic epitopes. The ability of our engineered antigens to induce neutralizing antibodies in guinea pigs validates the viability of our approach, and suggests it could apply to vaccine candidates still in the early stages of discovery to facilitate future translational development.

The reduction of product-related variants and increase in observed titers for all three antigens realized here enabled a uniform and simple purification of each antigen. Ultimately this simplicity aided the concomitant manufacturing of all three antigens required for a trivalent vaccine candidate. Specifically, reducing product-related variants by molecular engineering up front reduced the purification process to just two steps. Concomitant manufacturing of multiple serotypes, if performed consistently, could eliminate further steps in manufacturing like blending of antigens. Concomitant manufacturing could also significantly reduce the cost of manufacturing by minimizing the required facility time and space, as well as the number of tests and reviews needed for quality assessments and assurances [47].

## Conclusion

In this study, we have demonstrated that conservative molecular engineering can increase volumetric productivity, reduce unit operations, and enable concomitant manufacturing, all of which could reduce the cost of vaccine manufacturing [48]. A second implication of this study is that simple changes to a product sequence early in development can have a profound impact on the simplicity and scalability of the manufacturing process, potentially impacting further advancement of the product into the clinic and beyond. Separately, the accessibility of - omics data is exceptional for yeasts and other eukaryotic hosts, which enables further understanding of host biology for identifying rational host-dependent changes for product sequences [49]. This accessibility paired with the speed of cultivation and screening for expression suggests that yeast like *K. phaffii* are uniquely suited for the development of subunit vaccines. We believe our approach to molecular engineering demonstrates that subunit vaccines can be intentionally designed to conform to platform-like processes, achieving the same benefits in standardization seen with the manufacturing of monoclonal antibodies.

## Materials and methods

### Strains and cultivations

Wild-type *Komagataella phaffii* (NRRL Y-11430) was modified to express variants of P[4], P[6], or P[8] under control of the *AOX1* promoter using a commercial vector (pPICZ A, Thermo Fisher Scientific). Modified vectors containing markers for selection on G418 (Thermo Fisher Scientific) or Nourseothricin (Gold Biotechnology) were also used to generate a single strain expressing all three serotypes.

Strains for initial characterization were grown in 3 mL culture in 24-well deep well plates (25°C, 600 rpm) using complex medium (BMGY-Buffered Glycerol Complex Medium, Teknova) supplemented to 4% (v/v) glycerol. After 24h of biomass accumulation, cells were pelleted and resuspended in BMMY (Buffered Methanol Complex Medium, Teknova) containing 1.5% (v/v) methanol. Samples for RNA-seq and ribosome profiling were collected after 16h growth in BMMY.

Strains were additionally cultivated in 200 mL culture in 1 L shake flasks to generate material for non-clinical studies and in InSCyT[35] bioreactors for demonstration of end-to-end manufacturing. At both scales, cells were grown in rich defined medium[51] supplemented to 4% (v/v) glycerol for biomass accumulation or 5% (v/v) methanol for production. In the bioreactor, temperature, pH, and dissolved oxygen were maintained at 25°C, 6.5, and 25%, respectively.

### Transcriptome analysis

RNA was extracted and purified according to the Qiagen RNeasy kit (cat #74104) and RNA quality was analyzed to ensure RNA Quality Number >7. RNA libraries were prepared using the Roche KAPA HyperPrep kit and sequenced on an Illumina NextSeq to generate 40-nt paired-end reads.

Sequenced reads were aligned and quantified using STAR v2.5.3a[52] and RSEM v1.3.0[53]. Expression was visualized using *log_2_*(*Fragments per Kilobase of Transcript per Million Mapped Reads + 1*) values. Differential gene expression was analyzed using the DESeq2 package in R starting from gene integer counts and including log-fold-change shrinkage. Gene set enrichment analysis (GSEA) was performed in R using the fgsea package.[54] Gene expression data have been deposited in NCBI’s Gene Expression Omnibus and are accessible through GEO Series accession number GSE159325.

### Ribosome footprinting

Cell samples from cultures expressing P[4] or P[6] were transferred to a shake-flask containing 500 mL of pre-warmed fresh BMMY media to achieve an OD_600_ = 0.5. Cell harvest, cell lysis, and recovery of ribosome footprints were performed as described previously.[55] Footprints were dephosphorylated and cDNA libraries were prepared using the SMARTer smRNA-Seq Kit (Clontech), with an additional rRNA depletion step prior to PCR amplification, all as described previously.[55]

Libraries were sequenced on an Illumina NextSeq to generate 75-nt paired-end reads. The reverse reads were of low quality as expected from the library preparation kit and were not used in analysis. For the forward reads, the first 3 nucleotides and poly(A) sequences were removed from raw FASTQ reads using Cutadapt v.1.16[56] with options –u 3 -a A{100} –m 12. The trimmed reads were then aligned to a target consisting of *K. phaffii* rRNA sequences with bwa v0.7.16a. Unaligned reads were extracted with samtools v1.5 and FASTQ files were regenerated with bedtools v.2.26.0 for further analysis. Reads were then aligned to the wild-type *K. phaffii* genome plus the P[4] or P[6] expression cassettes using STAR v.2.5.3a[52] with the -- alignIntronMax 600 option. Secondary alignments were removed using samtools v.1.5 and the primary alignment bam files were used as input to plastid v.0.4.8.[57] Plastid was run according to the author’s documentation and the commands used are available in the file plastid_processing.sh, which can be found in the Github repository jrbrady / PlastidCommands. The read sizes considered ranged between 26 and 31 nt, and an offset of 13 nt was used for all lengths except 26 nt, for which a 12 nt offset was used. Ribosome profiling data have been deposited in NCBI’s Gene Expression Omnibus and are accessible through GEO Series accession number GSE159336.

### Experimental design

The goal of the animal study was to compare engineered antigens manufactured in *K. phaffii* to the original antigens manufactured in *E. coli*. We studied nine groups of five animals each, including a group for each of the original antigens (3), a group for each of the engineered antigens (3), one group of the original P[4] antigen manufactured in *K. phaffii*, one group receiving a placebo (adjuvant + buffer), and one group receiving trivalent rotavirus vaccine (Table S3).

### Immunization of guinea pigs

Groups of 5 guinea pigs (Hartley, Sisi Al strain) weighing 400-500g (National Institute of Agricultural Technology, INTA, laboratory animal production facility) were immunized intramuscularly three times at two-week intervals with 200 μl of aqueous vaccine containing 12 μg of the recombinant antigen (Table S3) adjuvanted with 2% Alhydrogel (Brenntag Biosector, Frederikssund, Denmark) (aluminum content 100 μg/dose). A group of animals immunized with a trivalent RVA vaccine formulated with a mix of HRVA Wa, DS1 and ST3 at 10^7^ FFU/dose in Alhydrogel was used as positive control. A group of 5 animals receiving buffer in Alhydrogel was used as negative control. Blood samples were collected prior to each immunization as well as at 7, 14, and 28 days after the third immunization. All guinea pig experiments were conducted in compliance with the guidelines of the Institutional Animal Welfare Committee of the National Institute of Agricultural Technology, INTA, CICUAE protocol # 55.2018.

### Neutralization and protein specific ELISA assays

Human rotavirus (HRVA) Wa GP[8], DS1 G2P[4] and ST3 G4P[6] adapted for cell culture were grown in MA104 cells using standard procedures. Viral titer was determined by cell culture immuno-fluorescence (CCIF) assay and expressed as FFU/ml as previously described.[58]

Virus neutralizing antibody titers (VNtAb) in serum samples were determined by a modified fluorescence focus reduction neutralization assay.[59] Serial two-fold dilutions of each serum sample were mixed with an equal volume of virus to a final concentration of 100 FFU. Serum-virus mixes were incubated at 37°C, 5% CO_2_ for 1h, and then transferred onto confluent monolayers of MA-104 cells in 96 well plates. Plates were incubated at 37°C, 5% CO_2_ for 48h. The monolayers were fixed with 70% acetone and the presence of residual virus in each well was evidenced by immuno-staining using a 2KD1 nanobody to rotavirus VP6 protein labeled with Alexafluor 488.[60] VNtAb of each serum sample is expressed as the inverse of the maximal dilution showing 60% fluorescence focus reduction.

Homotypic and heterotypic IgG antibody responses to vaccination were monitored using standard P2-VP8 antigen ELISA techniques. Individual serum samples were evaluated against the homologous antigen while the heterologous response was evaluated in serum pooled at every time point (T0 to T6). Thermo Fisher Maxisorp® high-binding 96-well ELISA plates were coated with each antigen diluted in carbonate buffer pH 9.6 for 24 hours at 4°C. Plates were washed with wash buffer (PBS with 0.05% TweenTM 20), blocked with 200 μL of blocking buffer (PBS with 10% Skim milk), and incubated for one to four hours at 37°C. Serum samples were diluted in serial four-fold dilutions, starting at 1:160, and 100 μL of each dilution was transferred to the assay plate, incubated for one hour at 37°C and washed as described above. Plates were incubated for one hour at 37°C with peroxidase labeled goat anti-guinea pig IgG antibody (Jackson) diluted 1:3000 in wash buffer. After four washes, 100 μL per well of detection reagent (H_2_O_2_, ABTS in citrate buffer, pH 5) was allowed to develop for 15 minutes in the dark at room temperature, and stopped with 50 μL of 5% SDS. Absorbance was measured at 405 nm with a SpectraMax® M2 microplate reader (Molecular Devices, LLC). The VNtAb of each serum sample was expressed as the inverse of the maximal dilution showing 60% fluorescence focus reduction. Serum that yielded no signal for any technique at 1:160 dilution was assigned a titer of 1:80.

### Protein purification

Protein purification for non-clinical studies and end-to-end manufacturing was carried out on the purification module of the InSCyT system as described previously[35]. All columns were equilibrated in the appropriate buffer prior to each run. Product-containing supernatant was adjusted to pH 4.5 using 100mM citric acid. The adjusted supernatant was loaded into a pre-packed CMM HyperCel column (1-mL or 5-mL) (Pall Corporation, Port Washington, NY), re-equilibrated with 20 mM sodium citrate pH 4.5, washed with 20 mM sodium phosphate pH 6.0, and eluted with 20 mM sodium phosphate pH 7.0, 100 mM NaCl. Eluate from column 1 above 15 mAU was flowed through a 1-mL pre-packed HyperCel STAR AX column (Pall Corporation, Port Washington, NY). Flow-through from column 2 above 15 mAU was collected.

### Analytical assays for product characterization

Biolayer inferometry was performed using the Octet Red96 with streptavidin (SA) biosensors (Pall ForteBio, Fremont, CA), which were hydrated for 15 min in kinetics buffer prior to each run. Kinetics buffer consisting of 1X PBS pH 7.2, 0.5% BSA, and 0.05% tween 20 was used for all dilutions, baseline, and disassociation steps. Monoclonal antibodies specific to P[4], P[6], and P[8], respectively, were biotinylated using the EZ-Link Sulfo-NHS-LC-biotinylation kit according to manufacturer instructions (Thermo Fisher Scientific) and used in the assay at concentrations of 1.25, 1.7, and 1 μg/mL, respectively. Triplicate samples were loaded in a 96-well black microplate (Greiner Bio-One) at starting concentrations of 2, 14, or 5 μg/mL, respectively. Seven 1:1 serial dilutions and a reference well of kinetics buffer were analyzed for each sample. Association and disassociation were measured at 1000 rpm for 300 and 600 sec, respectively. Binding affinity was calculated using the Octet Data Analysis software v10.0 (Pall ForteBio), using reference subtraction, baseline alignment, inter-step correction, Savitzky-Golay filtering, and a global 1:1 binding model.

Wet cell weight (WCW) was determined as described previously.[35] Sample concentrations were determined by absorbance at A280 nm. Sodium dodecyl sulfate polyacrylamide gel electrophoresis (SDS-PAGE) was carried out under reducing and nonreducing conditions using Novex 12% Tris-Glycine Gels or Novex 16% Tricine Gels (Thermo Fisher Scientific, Waltham, MA) according to the manufacturer’s recommended protocol and stained using Instant Blue Protein Stain (Thermo Fisher Scientific, Waltham, MA). Isoelectric focusing was carried out as described previously.[35] Samples were analyzed for host cell-protein content using the Pichia pastoris 1st generation HCP ELISA kit from Cygnus Technologies (Southport, NC) according to the manufacturer’s recommended protocol. Samples were analyzed for residual host-cell DNA using the Quant-iT dsDNA High-Sensitivity Assay Kit (Invitrogen) according to the manufacturer’s protocol except the standard curve was reduced to 0-20 ng.

High performance liquid chromatography (HPLC) analysis was performed on an Agilent 1260 HPLC system equipped with a diode array detector and controlled using OpenLab CDS software (Agilent Technologies, Santa Clara, CA). Antigen concentration was determined using a PLRP-S column (2.1 x 150 mm, 300Å, 3μm) operated at 0.5 mL/min and 60°C (Agilent Technologies, Santa Clara, CA). Buffer A was 0.1% (v/v) TFA in water and buffer B was 0.1% (v/v) TFA, 0.5% (v/v) water in ACN. A gradient of 37-42% B was performed over 30 minutes; total method run time was 45 minutes. Sample injection volumes were 10μL. Data analysis was completed using OpenLab CDS Data Analysis (Agilent Technologies, Santa Clara, CA).

Intact mass analysis was performed on a 6230B time of flight (TOF) LC-MS with a 1220 series HPLC (Agilent Technologies, Santa Clara, CA). Mobile phase A consisted of water with 0.1% formic acid, and mobile phase B was acetonitrile with 0.1% formic acid. About 20 pmol of each sample was injected, bound to a ZORBAX 300SB C3 column (Agilent Technologies), desalted, and subjected to electrospray ionization. The LC gradient consisted of 20-70% B over 1 min at a flow rate of 1.5 mL/min. Elution of proteins was monitored using the absorbance signal at 214 nm. 50 μL of isopropanol was injected after each sample to control sample carryover. The typical electrospray ionization parameters consisted of: 290°C gas temperature, 4000 V Vcap, 2000 V nozzle, and 275 V fragmentor voltage. Mass spectra were collected from 700-2800 m/z at a scan rate of 1 spectra/sec. MS spectra were processed using MassHunter Qualitative Analysis software (v B.07.00, Agilent Technologies) with deconvolution range of 10-50 kDa, using 1 Da mass step.

## Declarations

### Ethics approval and consent to participate

Animal studies were reviewed and approved by The Institutional Committee for the Care and Use of Experimental Animals at INTA CICVyA, Argentina, evaluation request number 55/2018, “Immunogenic quality control of subunit vaccines for human rotavirus in guinea pigs.”

## Consent for publication

Not applicable.

## Availability of data and materials

Sequencing data have been deposited in NCBI’s Gene Expression Omnibus (GEO) and are accessible through GEO Series accession number GSE159338.

## Competing interests

L.E.C., K.R.L., and J.C.L. have filed patents related to the InSCyT system and methods. K.R.L. and J.C.L. are co-founders and consultants to Sunflower Therapeutics PBC.

## Funding

This work was funded by the Bill and Melinda Gates Foundation (Investment ID OPP1154682). This study was also supported in part by the Koch Institute Support (core) grant P30-CA14051 from the National Cancer Institute. J.R.B. and N.C.D were partially supported by a NIGMS/MIT Biotechnology Training Program Fellowship (NIH contract no. 2T32GM008334-26). The content is solely the responsibility of the authors and does not necessarily represent the official views of the NCI, NIH, or the Bill & Melinda Gates Foundation.

## Authors’ contributions

N.C.D., J.R.B., K.R.L., and J.C.L. conceived and planned experiments. N.C.D., J.R.B., M.K.T., and D.L.K. generated and characterized yeast strains. J.R.B. and C.A.W. conducted transcriptomic and ribosome profiling experiments. L.E.C. designed and performed protein purifications. A.M.B. and A.B. operated bioreactors. S.R.A. performed HPLC assays. K.K. and J.M.H. performed analytical characterization. M.B., C.V., and V.P. performed animal studies. N.C.D., J.R.B., K.R.L., and J.C.L. wrote the manuscript. J.C.L., K.R.L., V.P., D.B.V., S.B.J., and T.M. designed the experimental strategy and supervised analysis. All authors reviewed the manuscript.

## Acknowledgements

The authors thank Danielle Camp for program support and alignment, the Koch Institute Integrated Genomics and Bioinformatics Core for technical support, James Taggart for support with ribosome profiling, Nishant Sawant for assistance in sample preparation for mass spec analysis, Lucia Rocha for help with ELISAs, cell culture, and VN assays, Vanesa Franco, Jose Vallejo, and Ezequiel Rivarola for care of guinea pigs, and Andres Wigdorovitz for helpful conversation. The authors acknowledge PATH for providing reference NRRV antigens and related immunological mAb reagents for this study.

## Supplemental materials

**Fig. S1.**
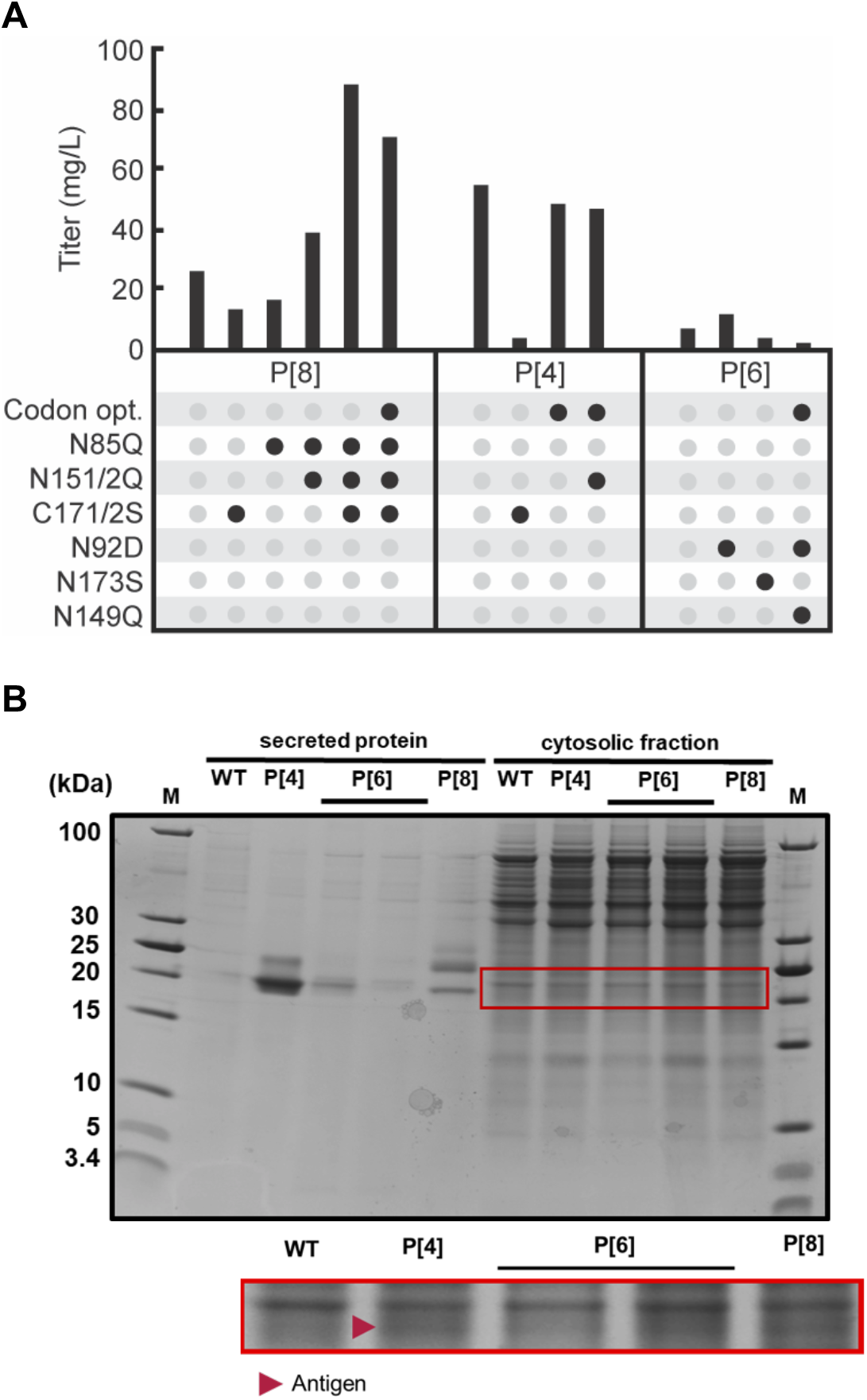
Titers of antigen expression. A) Secreted titers of each NRRV antigen variant produced in 3 mL, plate-based cultures. Titer was calculated using SDS-PAGE and densitometry, with comparison to a known standard. B) SDS-PAGE of supernatant and intracellular lysates from expression of original serotypes. No significant amount of P[6] antigen was detected in cell lysates.

**Fig. S2.**
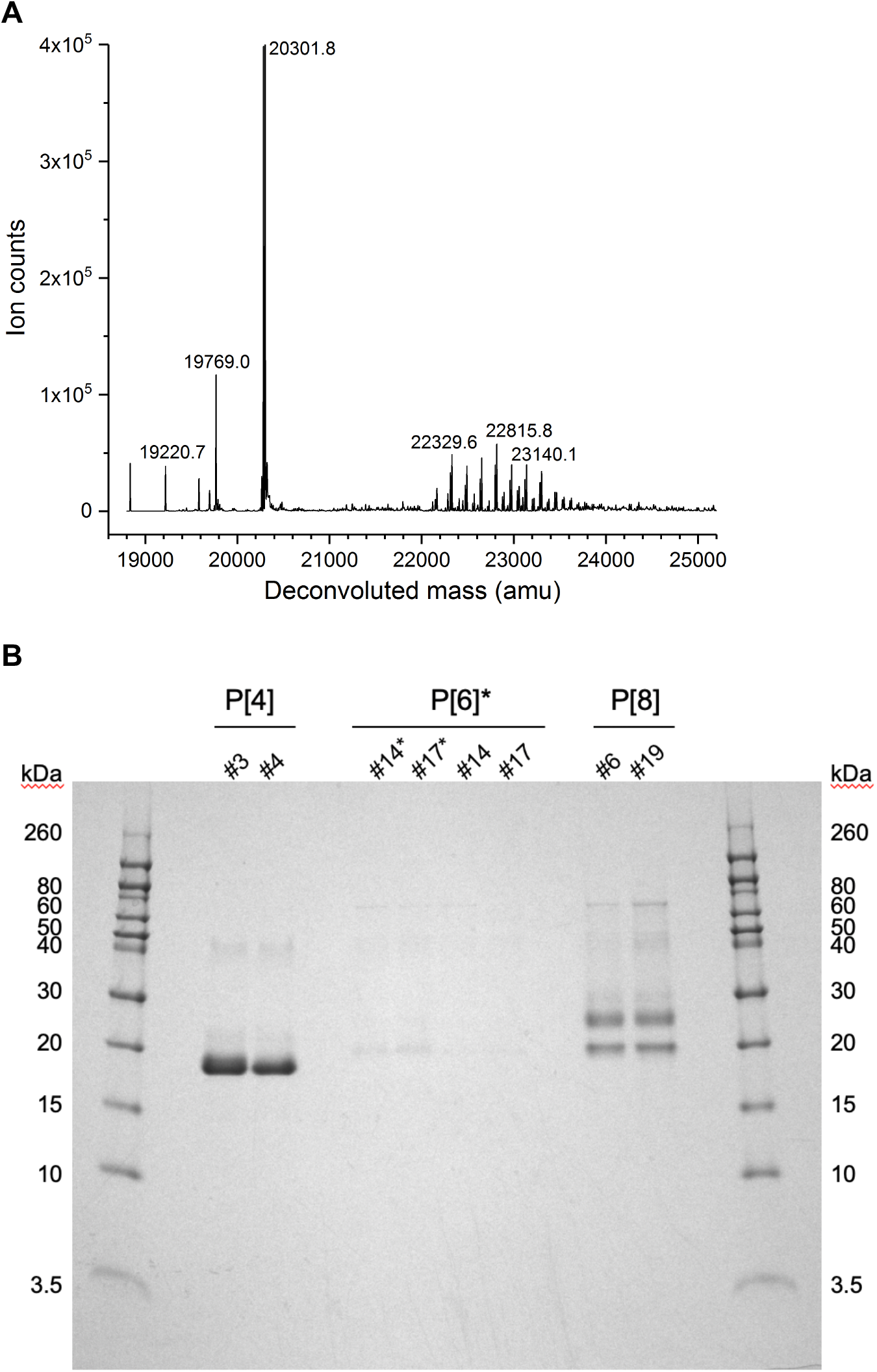
Quality analyses of antigens. A) Mass spectrum of P[8] from intact LCMS showing putative full-length, aglycosylated P[8] (~20.3 kDa) and high-mannose variants of P[8] (>22 kDa). B) Non-reduced SDS-PAGE of P[4], P[6], and P[8] supernatants. P[6] samples with an asterisk are concentrated 5x. Dimer and higher aggregates are visible in P[4] and P[8] lanes.

**Fig. S3.**
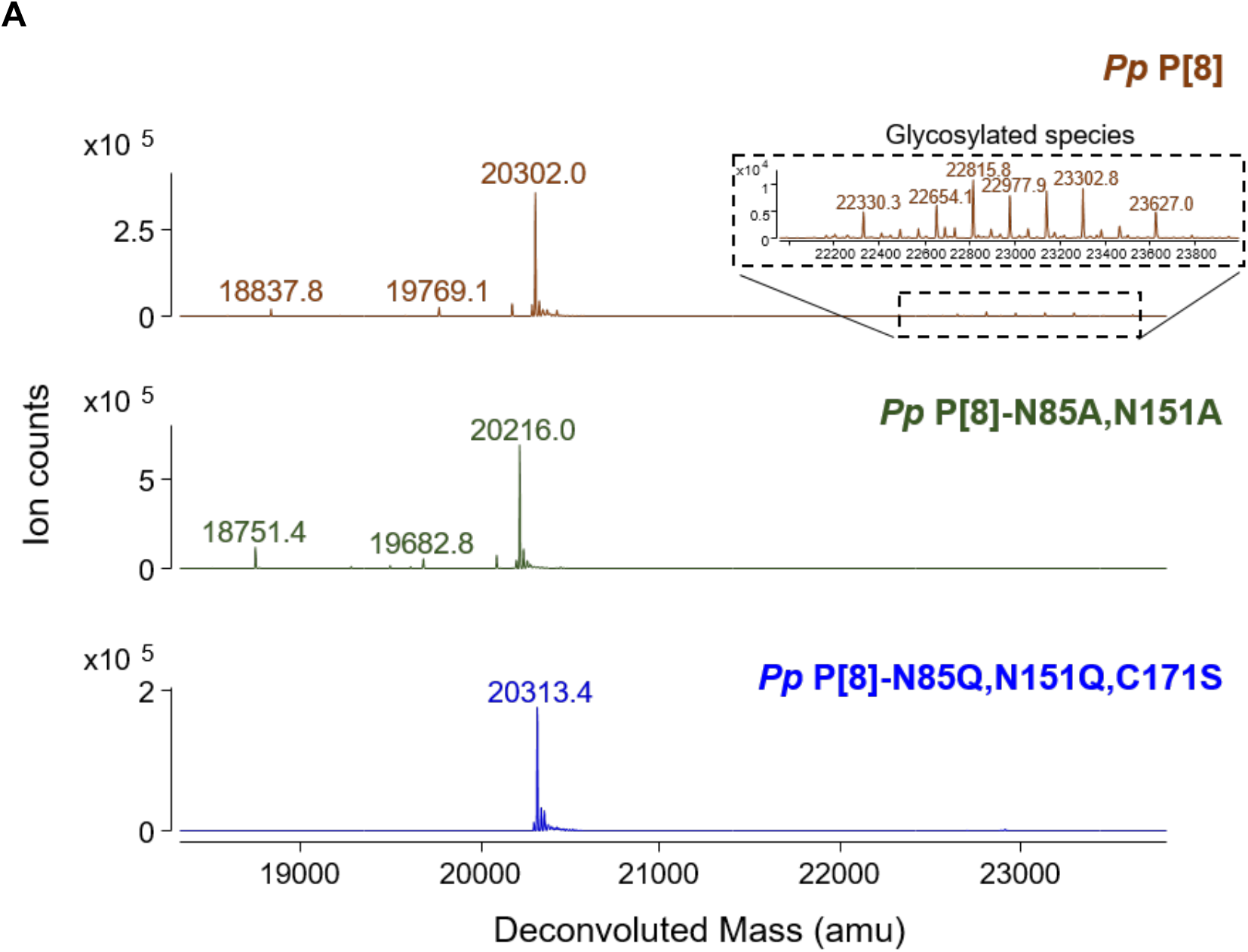
Glycosylation removal in P[8]. A) Mass spectra of P[8] variants, indicating the removal of hypermannosylated product variants with the introduction of N85Q and N151Q mutations.

**Fig. S4.**
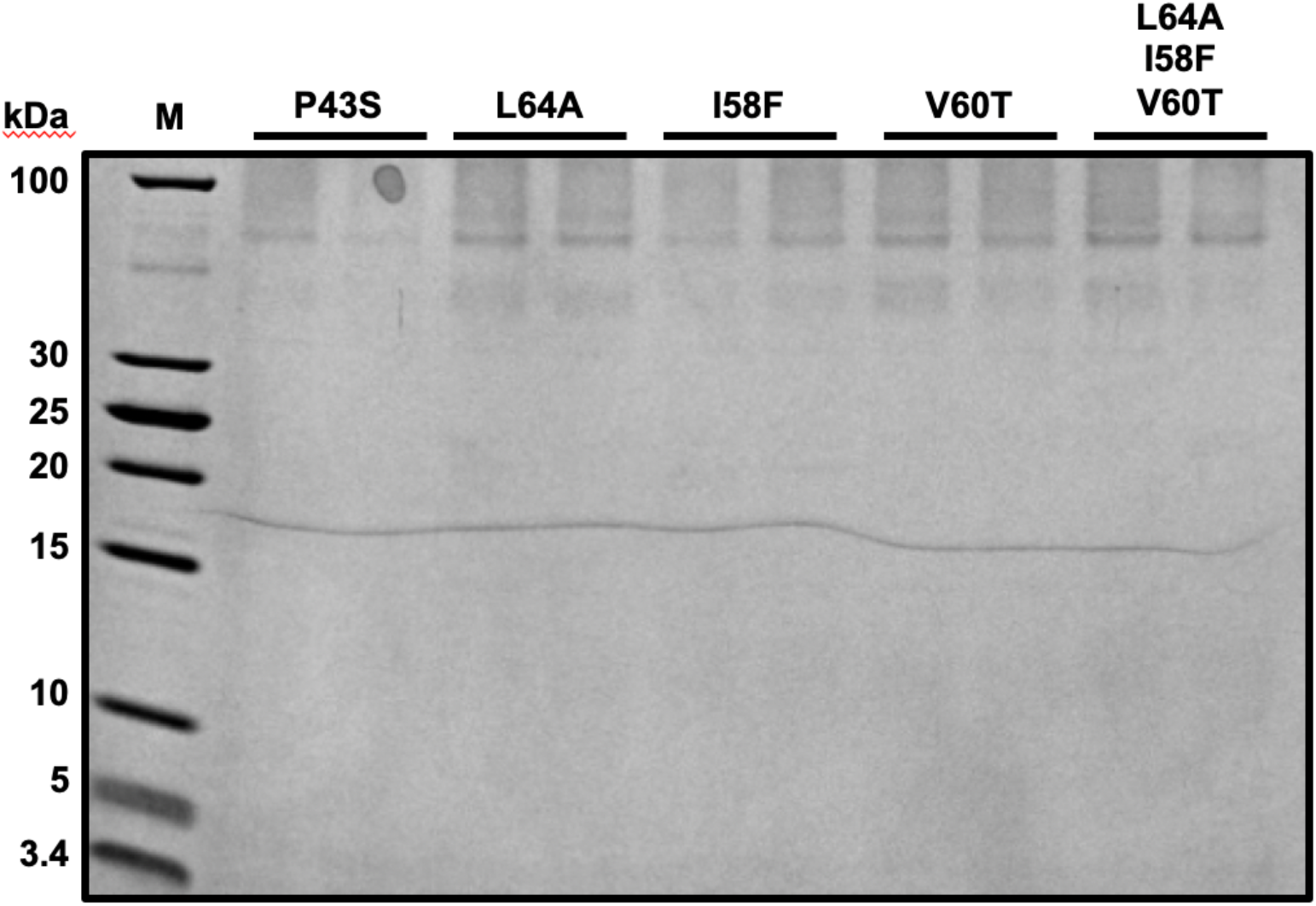
Hydrophobic region mutations in P[6]. A) SDS-PAGE of sequence variants of P[6] with reduced hydrophobicity. Expected product size is ~20 kD.

**Fig. S5.**
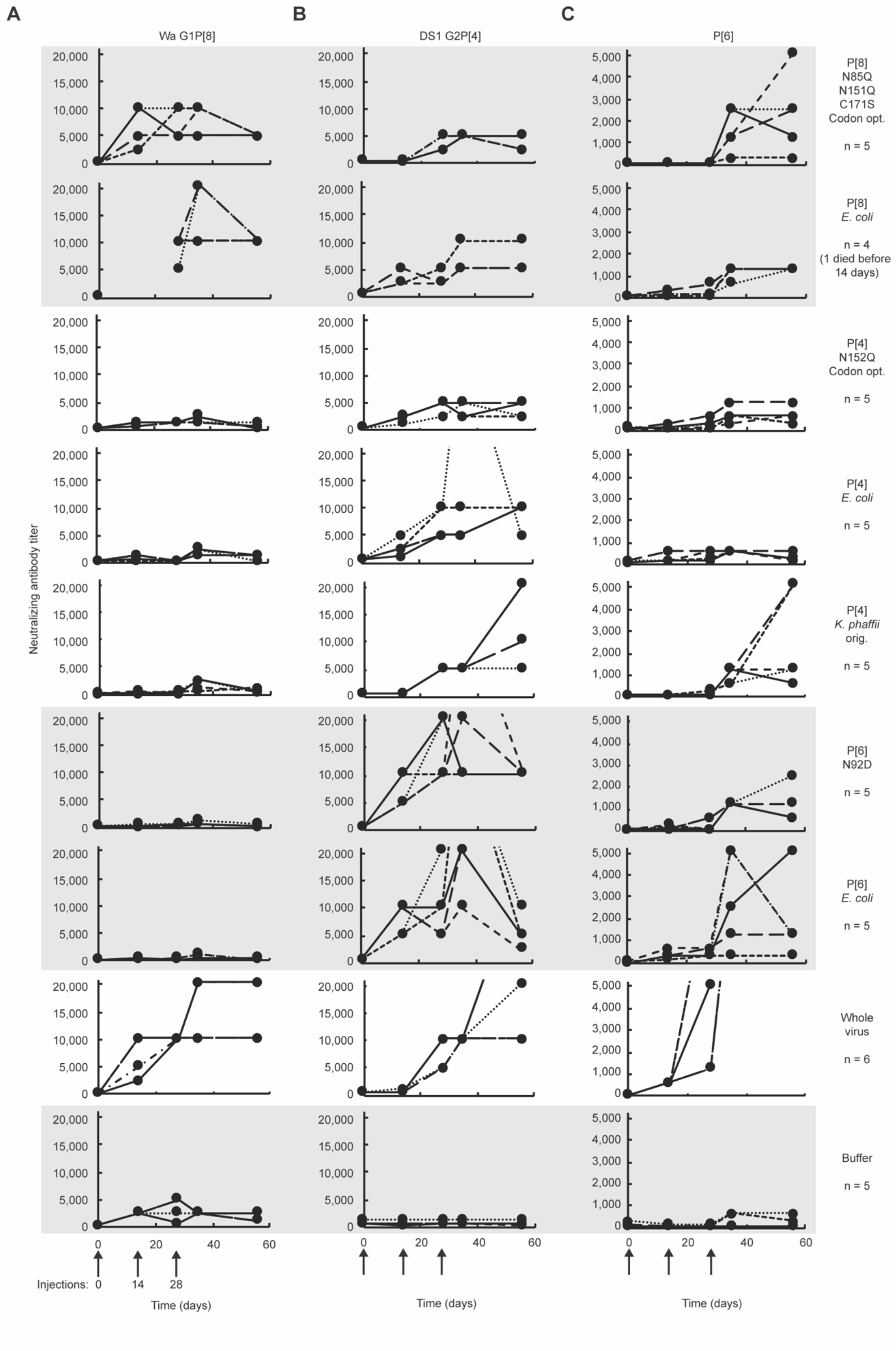
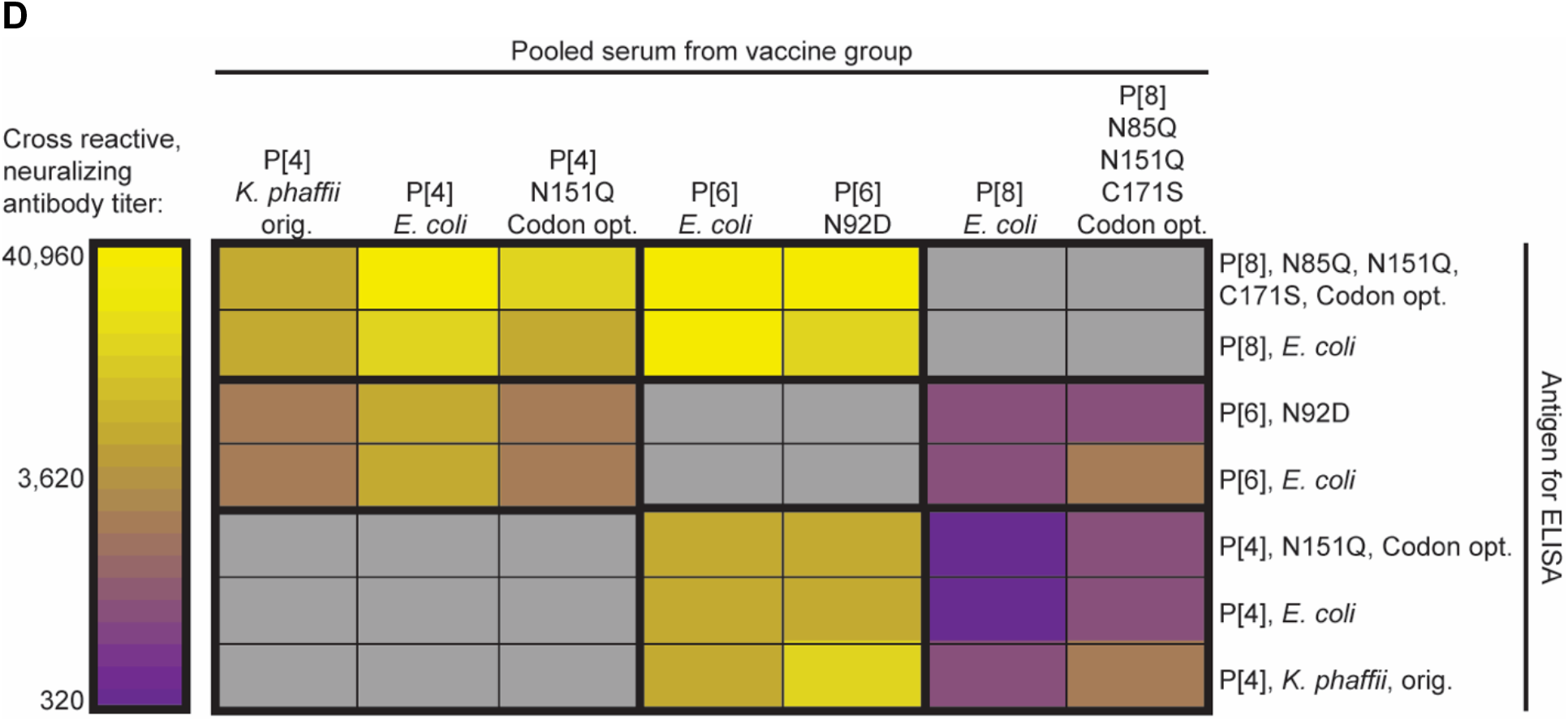
Animal study results. Titer over time of neutralizing antibodies generated in guinea pigs against A) P[8], B) P[4], and C) P[6] virus strains. Each line represents one animal, vaccinated with the antigen indicated at right. Animals were immunized at 0, 14, and 28 days for a total of three doses. Lines extending above the chart limits indicate a titer of 40,960, or 10,240 for ST3 G4P[6]. D) Cross reactivity of serum across all seven antigens by ELISA for IgG antibodies. Serum from all animals in each group was pooled.

**Fig. S6.**
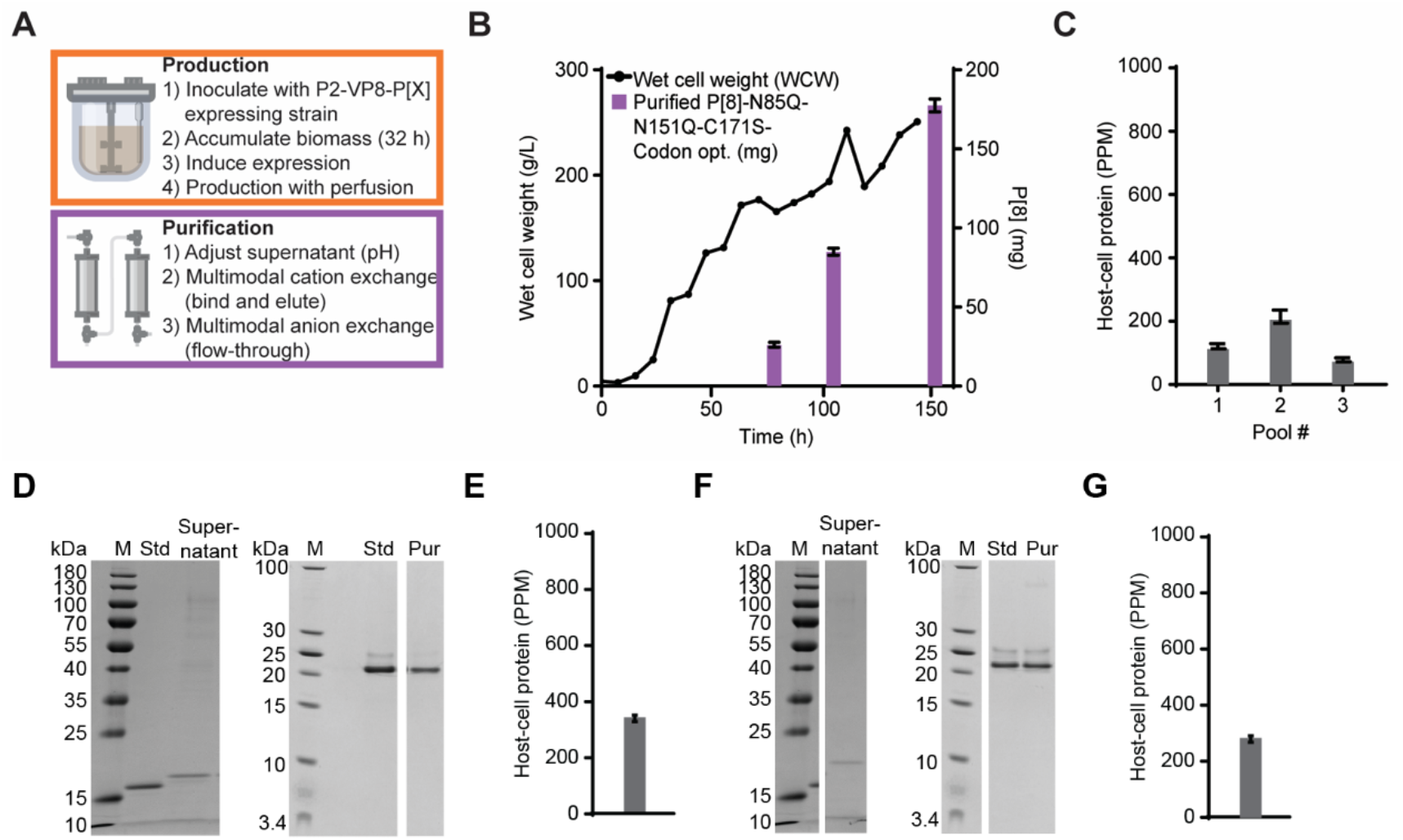
Process development of individual antigens. A) Schematic and B,C) process development data for an end-to-end production run of engineered P[8]. D-G) Purification development of engineered versions of P[4] (D,E), and P[6] (F,G). Cells were cultured in shake flasks, and supernatant was purified using the same process as engineered P[8].

**Fig. S7.**
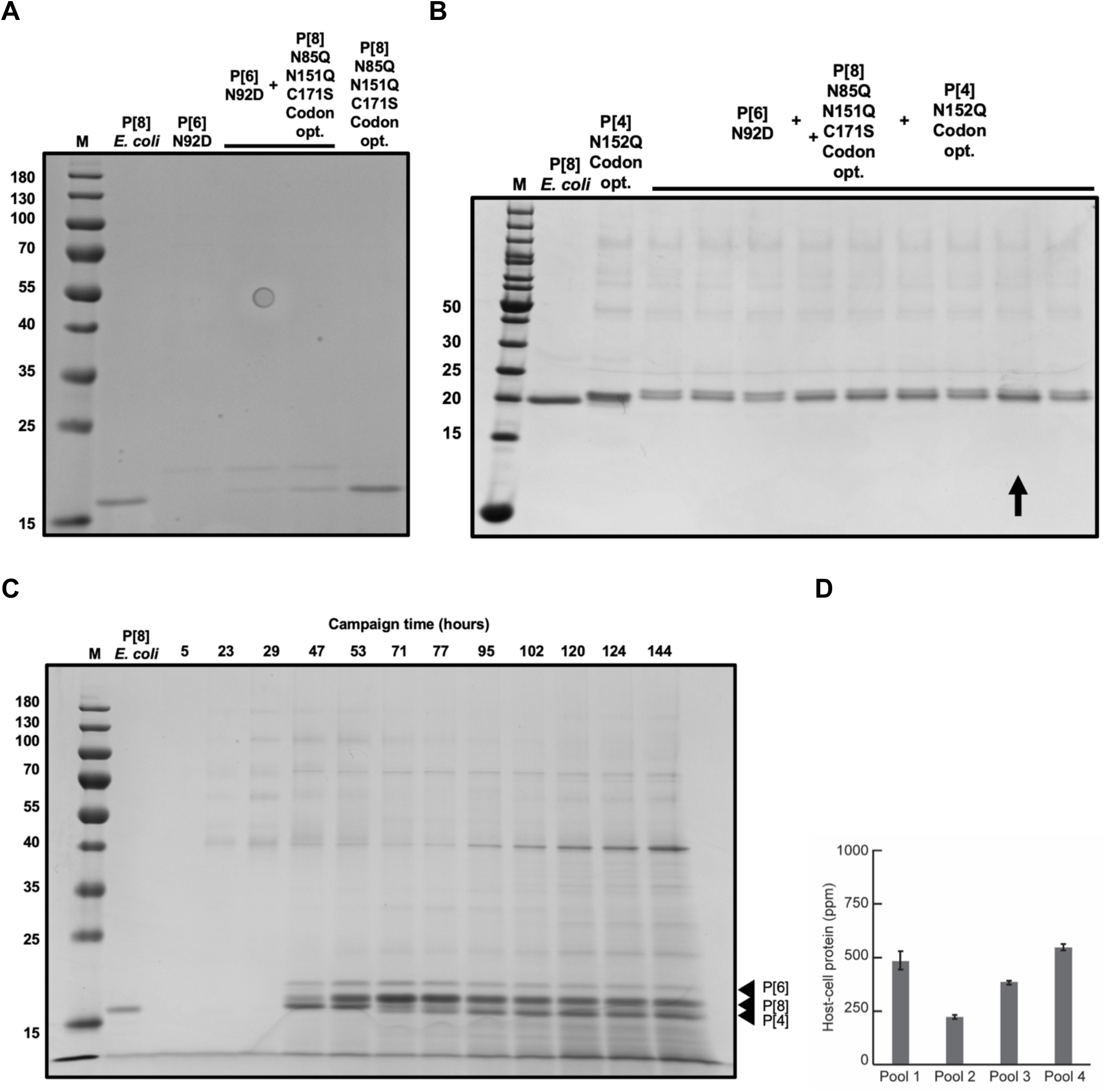
Expression of all three NRRV antigens. A) SDS-PAGE of supernatant of two clones that express engineered P[8] in a background strain expressing engineered P[6]. B) SDS-PAGE of supernatant of cells that express all three antigens. Cells were cultivated at 3 mL plate scale. The strain marked with an arrow was carried forward to reactor scale. C) SDS-PAGE of bioreactor samples across the campaign. All three engineered antigens are visible by SDS-PAGE. D) Concentration of host cell protein in purified product pools, as measured by host cell protein ELISA for *K. phaffii*.

**Table S1.** Accessibility scores of potential *N*-linked glycosylation sites in P[8], identified with NetNGlyc.

| Index | N-X-S/T | Accessibility score |
| --- | --- | --- |
| 53 | N-N-S | 0.2 |
| 85 | N-V-S | 0.7 |
| 88 | N-D-S | 0.2 |
| 151 | N-I-S | 0.9 |

**Table S2.** Normalized enrichment scores for gene sets significantly enriched (p-adj <0.05) in genes differentially expressed between serotype-expressing strains.

| **Gene ontology - biological process** | **P[6] vs. Null** | **P[6] vs. P[4]** | **P[6] vs. P[8]** |
| --- | --- | --- | --- |
| cytoplasmic translation | 2.19 | 2.52 | 2.19 |
| translocation | 1.92 | 1.81 |  |
| protein glycosylation | 2.13 | 2.22 | 2.02 |
| cell wall organization or biogenesis | 2.38 | 2.18 | 2.28 |
| DNA replication | 1.78 | 1.96 | 2.20 |
| carbohydrate metabolic process | 1.68 | 1.88 | 2.18 |
| DNA repair |  | 1.79 | 2.16 |
| telomere organization | 1.89 | 1.78 | 1.95 |
| cell budding | 1.80 | 1.72 | 1.97 |
| DNA recombination |  | 1.66 | 2.04 |
| cellular response to DNA damage stimulus |  | 1.63 | 2.02 |
| signaling | 1.73 | 1.55 | 1.65 |
| cytoskeleton organization | 1.68 | 1.51 | 2.11 |
| Golgi vesicle transport | 1.65 | 1.51 | 1.39 |
| cell morphogenesis |  |  | 1.84 |
| chromatin organization |  |  | 1.50 |
| chromosome segregation |  |  | 1.97 |
| cofactor metabolic process |  |  | 1.63 |
| conjugation | 1.78 |  | 1.80 |
| cytokinesis |  |  | 1.75 |
| generation of precursor metabolites | -1.74 |  | 1.82 |
| meiotic cell cycle |  |  | 1.89 |
| mitochondrial translation |  |  | -2.01 |
| mitotic cell cycle | 1.62 |  | 1.93 |
| monocarboxylic acid metabolic process |  |  | 1.61 |
| mRNA processing |  |  | -1.61 |
| small molecule metabolic process |  |  | 2.06 |
| organelle fission |  |  | 1.96 |
| protein folding |  |  | -2.26 |
| regulation of cell cycle |  |  | 1.70 |
| regulation of protein modification |  |  | 1.57 |
| regulation of transport | 1.68 |  |  |
| ribosomal large subunit biogenesis |  |  | -1.52 |
| ribosomal small subunit biogenesis | 1.81 |  | -1.72 |
| RNA modification |  |  | -2.12 |
| RNA splicing |  |  | -1.63 |
| rRNA processing | 1.70 |  | -2.10 |
| sporulation |  |  | 1.59 |
| transposition |  |  | 1.69 |
| tRNA processing |  |  | -1.77 |
| vitamin metabolic process | -2.41 |  |  |

**Table S3.** Animal study groups.

| Host | Antigen | N | Notes |
| --- | --- | --- | --- |
| E. coli | P[8] | 5 | 1 animal died |
| E. coli | P[6] | 5 |  |
| E. coli | P[4] | 5 |  |
| K. phaffii | P[8]-N85Q-N151Q-C171S-Codon opt. | 5 |  |
| K. phaffii | P[6]-N92D | 5 |  |
| K. phaffii | P[4]-N152Q-Codon opt. | 5 |  |
| K. phaffii | P[4] | 5 |  |
| --- | RVA trivalent | 6 |  |
| --- | Buffer | 5 |  |

